# Spatial single cell transcriptomic analysis of a novel DICER1 Syndrome GEMM informs the cellular origin and developmental hierarchy of associated sarcomas

**DOI:** 10.1101/2024.11.30.624592

**Authors:** Felix K.F. Kommoss, Joyce Zhang, Branden J. Lynch, Shary Yuting Chen, Janine Senz, Yana Moscovitz, Lesley Ann Hill, Wilder Scott, Jonathan Bush, Kenneth S. Chen, Andreas von Deimling, William D. Foulkes, Gregg Morin, T. Michael Underhill, Yemin Wang, David G. Huntsman

## Abstract

DICER1 syndrome predisposes children and young adults to tumor development across various organs. Many of these cancers are sarcomas, which uniquely express the RNase IIIb domain-deficient form of DICER1 and exhibit consistent histological and molecular similarities regardless of their anatomical origins. To uncover their cellular origin and developmental hierarchy, we established a lineage-traceable genetically engineered mouse model that allows for precise activation of *Dicer1* mutations in Hic1^+^ mesenchymal stromal cells. This model resulted in the development of renal tumors closely mirroring human DICER1 sarcoma histologically and molecularly. Single-cell transcriptomics coupled with targeted spatial gene expression analysis revealed a Hic1^+^ progenitor population marked by *Pdgfra*, *Dpt*, and *Mfap4,* corresponding to universal fibroblasts of steady-state kidneys. These fibroblastic progenitors exhibit the capacity to undergo rhabdomyoblastic differentiation or transition to highly proliferative anaplastic sarcoma. Investigation of patient samples identified analogous cell states. This study uncovers a fibroblastic origin for DICER1 sarcoma and provides a faithful model for mechanistic investigation and therapeutic development for tumors within the rhabdomyosarcoma spectrum.

## Introduction

DICER1 tumor susceptibility syndrome (DICER1 syndrome) is caused by germline pathogenic variants (PVs) in *DICER1* and predisposes children and young adults to cancer^1,2^. This includes several tumor types which arise at distinct anatomical locations, including the lungs, the kidney, the gynecologic tract, and the central nervous system (CNS), among other rarer sites^1–3^. Germline *DICER1* PVs typically inactivate the affected allele^4^. In DICER1 syndrome, tumor development is associated with the occurrence of a secondary missense mutations within the catalytic RNase IIIb domain of the second *DICER1* allele^5^. This compound heterozygous state of *DICER1* leads to impaired processing of pre-miRNAs and subsequent loss of mature miRNAs derived from their 5p-strands (5p-miRNA)^5–7^.

Many of the cancers associated with DICER1 syndrome represent clinically aggressive sarcomas. Although arising in different anatomical sites, it has long been noted that some of these neoplasms share key histomorphological features^8^. This includes subepithelial condensation of malignant mesenchyme (cambium layer), as well as frequent rhabdomyoblastic differentiation, resembling rhabdomyosarcoma (RMS). This overlap has raised important questions about whether these tumors arising at different sites constitute a common tumor entity. Recently we addressed this hypothesis by studying a large series of DICER1 syndrome associated cancers^9^. Our study identified shared clinicopathological, genetic and epigenetic (DNA methylation) features in a distinct group of DICER1-associated mesenchymal neoplasms. This group encompasses well established tumor entities and comprises three clinically meaningful classes termed “low-grade mesenchymal tumor with DICER1 alteration” (LGMT DICER1), “sarcoma with DICER1 alteration” (SARC DICER1), and primary intracranial sarcoma with DICER1 alteration (PIS DICER1). While our findings suggest a common cellular origin for DICER1 sarcoma, the underlying developmental biology and processes involved in DICER1-mediated tumor cell transformation remains largely unknown.

Genetically engineered mouse models (GEMM) are powerful tools for studying tumor development in a cell type specific manner. We have recently developed the first conditional *Dicer1* RNase IIIb mutant allele, from which we also developed the first-ever GEMM of DICER1 syndrome that recapitulates the compound heterozygous *Dicer1* mutations seen in most DICER1 syndrome-associated cancers^10^. In this model mutations were induced through an anti-Mullerian hormone receptor type 2 (Amhr2)-Cre driver. This led to activation of mutations in stromal cells throughout the Mullerian tract and adenosarcomas, a cancer type well described in DICER1 syndrome^11^. Subsequently, Oikawa et al. developed an Albumin-driven Cre with a human *DICER1* RNase IIIb hotspot mutation (G1809R) that developed epithelial lesions of the liver^12^. However, these and other models do not allow for controlling mutation activation and lineage tracing, significantly limiting their application to study tumor development^13,14^.

To further expand our existing model, control the timing of mutation activation, and enable lineage-tracing, we crossed our Dicer1 tool strains with our recently generated traceable CreERT2 mouse line-Hic1CreERT2, which expresses Cre in mesenchymal stromal cells (MSCs) throughout adult tissues upon tamoxifen treatment^15^. This resulted in the development of renal sarcomas that closely resemble cancers seen in the human disease. We leveraged single-cell whole transcriptomics (scRNA-seq) and a targeted spatial transcriptomics platform to uncover the putative cellular origin and developmental trajectory of Dicer1 sarcomagenesis in murine tumors which were compared to human disease.

## Results

### A lineage-traceable and genetically true model system of Dicer1 sarcoma

We utilized a tamoxifen-inducible Cre, driven by the Hypermethylated in Cancer 1 (Hic1) promoter, which marks quiescent mesenchymal progenitors in most vascularized tissues^15^. In parallel, the (Rosa26)^LSL-tdtomato^ strain permanently labeled cells with tdTomato fluorescence^16^. This led to the derivation of Hic1^CreERT2^;Dicer1^fl/fl-D1693N^;(Rosa26)^LSL-tdTomato^ mice (referred to as HDT hereafter, Fig. 1a). Activation of Dicer1 mutations postnatally, regardless of sex (Extended Data Fig. 1a), resulted in the development of renal tumors with 81.5% penetrance and a median latency of 378 days (range of 8-18 months) (Fig. 1b, Supplementary Table 1). Different control genotypes did not lead to development of renal tumors and did not differ in overall survival (Extended Data Figs. 1b-f). There was a moderate linear relationship between tamoxifen injection time and latency (Fig. 1c).

**Fig. 1:**
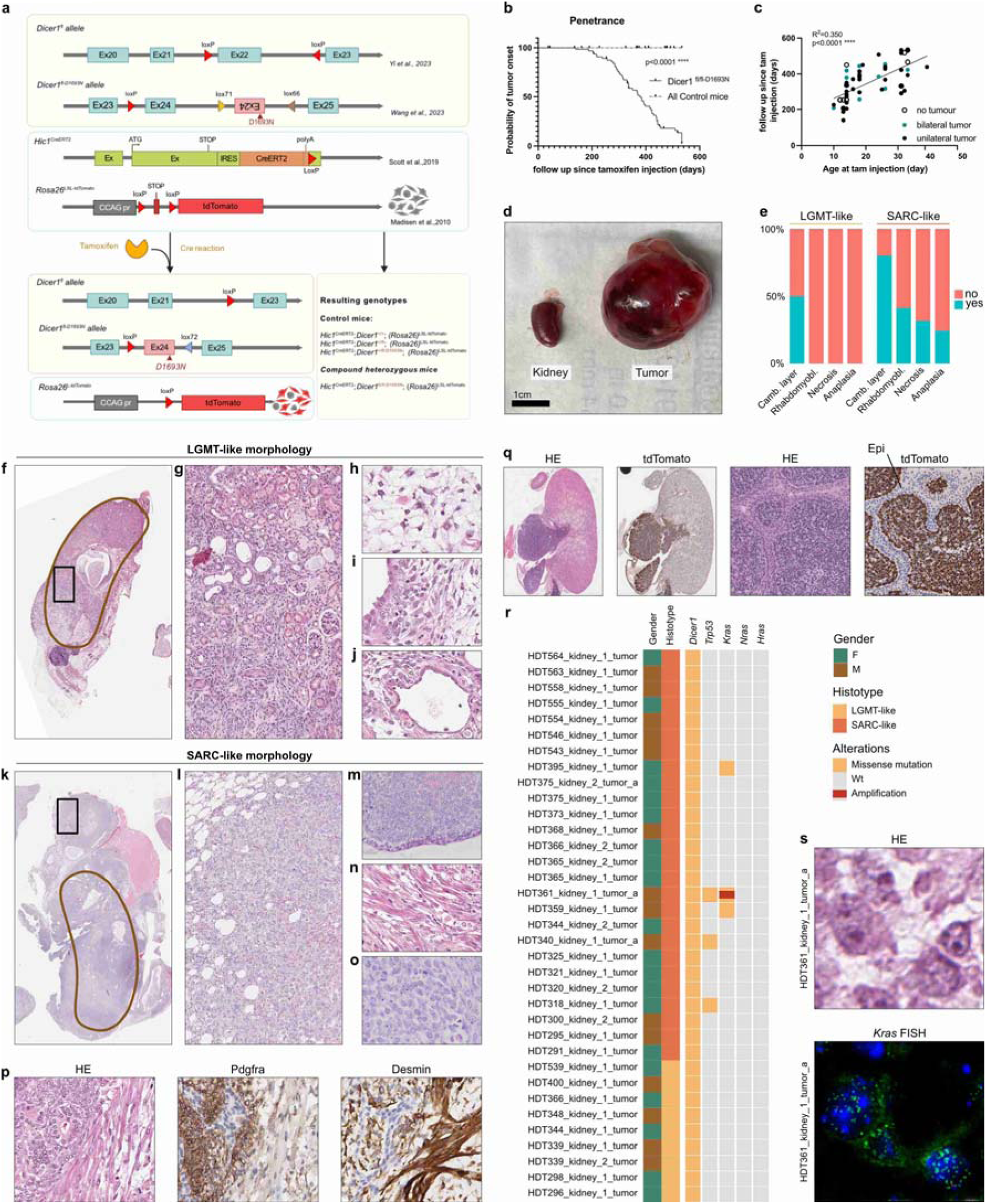
Breeding strategy and penetrance, histological and genomic characterization of a lineage-traceable and genetically true model system of Dicer1 sarcoma. a, Detailed description of HDT breeding strategy utilizing 4 existing alleles of Dicer1^fl^, Dicer1^fl-D1693N^, Hic1^CreERT2^, Rosa26^LSL-Tomato^. In HDT mice, tamoxifen injection leads to tdTomato expression and allows for lineage-tracing. Created with content from BioRender.com. b, Penetrance of tumor development in all control mice strains and HDT (Dicer1^fl/fl-D1693N^) with median latency of 378 days in HDT mice (range of 8-18 months). c, Correlation of follow-up days since tamoxifen injection and age at tamoxifen injection (tamoxifen injection day - date of birth). The linear relationship is fit with both unilateral and bilateral tumors using linear regression with Wald test. d, Comparison of a renal mass to a normal kidney from a HDT mouse. e-o, histological features of renal tumors: small lesion confined to the renal parenchyma showing features akin to human low-grade mesenchymal tumours (LGMT) with DICER1 alteration (f, g), including hypocellular mesenchyme (h) and focal condensation adjacent to cystically dilated renal ducts (i, j). Large lesion overgrowing the renal parenchyma showing features akin to human sarcoma (SARC) with DICER1 alteration (k), including sarcomatous mesenchyme (l), subepithelial aggregation of tumor cells (cambium layer, m), prominent rhabdomyoblastic differentiation (n), and anaplasia (o). p, Tumors stain for Pdgfra and Desmin, with Pdgfra marking areas of primitive tumor cells and Desmin highlighting regions of rhabdomyoblastic differentiation. q, While the mesenchyme of tumors is diffusely positive for tdTomato, non-neoplastic lining epithelium is negative for tdTomato throughout. q, Summary of mutations identified from targeted sequencing, including *Dicer1* (Asp1693Asn), *Trp53* (Lys162Arg, Arg172His, and Phe110Val), and *Kras* (Gln61His and Gly12Arg) of 35 HDT tumor samples, with clinical and histological information included as an oncoplot to the side. No mutations were identified in *Nras* or *Hras*. Histotype: Tumors with LGMT-like morphology, LGMT-like; tumors with SARC-like morphology, SARC-like.

Macroscopically, tumors originated from the kidney, presenting as a continuum that ranged from small tumors which maintain the integrity of the kidney structure to large masses that had completely overgrown the kidney (Fig. 1d). Microscopically, renal tumors closely mirrored human DICER1-associated mesenchymal neoplasms with tumor exhibiting a continuum from LGMT-like features to frank sarcomatous tumors showing features akin to SARC DICER1/PIS DICER1 (Fig. 1e, Supplementary Table 2). Notably, this continuum included histological features typically associated with cystic nephroma, which is classified under the LGMT DICER1 group, as well as anaplastic sarcoma of the kidney, categorized within the SARC DICER1 class^9^. In detail, this included small tumors, typically limited to the kidney’s parenchyma (Figs. 1f and g), exhibiting hypocellular and primitive mesenchyme (Fig. 1h) with focal condensation adjacent to cystically dilated renal ducts (Figs. 1i and j). Conversely, large tumors that typically showed sarcomatous overgrowth of neighboring tissue consisted of sarcomatous mesenchyme with a pattern-less or fascicular growth (Figs. 1k and l). In these tumors, subepithelial aggregation of tumor cells underneath transitional cell epithelium (bona fide cambium layer) (Fig. 1m), prominent rhabdomyoblastic differentiation (Fig. 1n), and prominent anaplasia (Fig. 1o), often accompanied with necrosis, was noted. In line with human disease, tumors stained for Pdgfra and Desmin^2,17^, with Pdgfra marking areas of primitive tumor cells and Desmin highlighting regions of rhabdomyoblastic differentiation (Fig. 1p). While the mesenchyme of all tumors was diffusely positive for lineage-tracing marker tdTomato, in contrast, associated lining epithelium was negative for tdTomato throughout (Fig. 1q), indicating that it was not developed from the neoplastic lineage. Targeted sequencing of HDT tumors confirmed the expected *Dicer1* RNase IIIb (D1693N) mutation in all sequenced samples and revealed recurrent hotspot mutations in *Kras* and *Trp53* which are commonly seen in human DICER1 sarcoma^9,18–21^ in 4/35 and 3/35 tumors, respectively (Fig. 1r, Supplementary Table 2). Furthermore, qPCR for *Kras* identified an amplification in one tumor with a concurrent *Kras* missense mutation, which was confirmed by fluorescence in-situ hybridization (FISH) (Fig. 1s). Notably, these mutations were only identified in neoplasms with SARC-like differentiation. Overall, these results demonstrate that renal neoplasms from our HDT GEMM develop from the Hic1+ lineage and recapitulate the tumor development continuum of human DICER1 syndrome-associated mesenchymal neoplasms.

### Organization of the Hic1^+^ mesenchymal stromal cell lineage in murine steady state kidney

In skeletal muscle, heart, and skin, Hic1 effectively identifies various MSC populations, including fibroblasts and pericytes, alongside rare perineural cells^15,22,23^. However, the characteristics of the Hic1^+^ MSC lineage in kidney remains elusive. To establish the steady-state composition of the renal Hic1+ MSC niche (Extended Data Fig. 2), we analyzed renal tissue from Dicer1^+/+^, Dicer1^+/fl^, and Dicer1^+/fl-D1693N^ control mice using scRNA-seq (*n*=3, PNW 4-32) and spatial transcriptomics (10X Genomics, Xenium, 379 genes; *n*=4, PNW 81) (Fig. 2a, Supplementary Table 3 and 4). ScRNA-seq identified multiple clusters correlating to physiological kidney cell composition (Extended Data Figs. 3a-d, Supplementary Table 5)^24–26^. Focused analysis of *tdTomato*^+^ MSCs identified 6 clusters (Extended Data Figs. 3e, Supplementary Table 2) that were grouped into 5 cell types based on marker expression (Fig. 2b-d, Extended Data Figs. 3f, Supplementary Table 6). This included a cluster of mural cells (Mural, *Myh11*^+^, *Acta2*^+^, *Tagln*^+^), and two types of *Pdgfra*^+^ and *Dpt*^+^ fibroblasts, one characterized by *Pi16^high^*and *Ly6a^high^* (Uni Fibro *Pi16*^high^) and the other by *Col15a1^high^* and *Mfap4^high^* (Uni Fibro *Col15a1*^high^). We furthermore identified two clusters of *Pdgfra*^+^ cells that were largely negative for *Dpt*, *Pi16* and *Col15a1*. One (Fibro) showed a signature resembling kidney interstitial fibroblasts and mesangial cells (*Itga8*+, *Robo2*+, *Ptn*+), while the other (Cycle) was enriched for cycling markers (*Cenpf*^+^, *Top2a*^+^, *Mki67*^+^). Notably, Uni Fibro *Pi16*^high^ and Uni Fibro *Col15a1*^high^ were more abundant in juvenile mice compared to adults (Extended Data Fig. 3g).

**Fig. 2:**
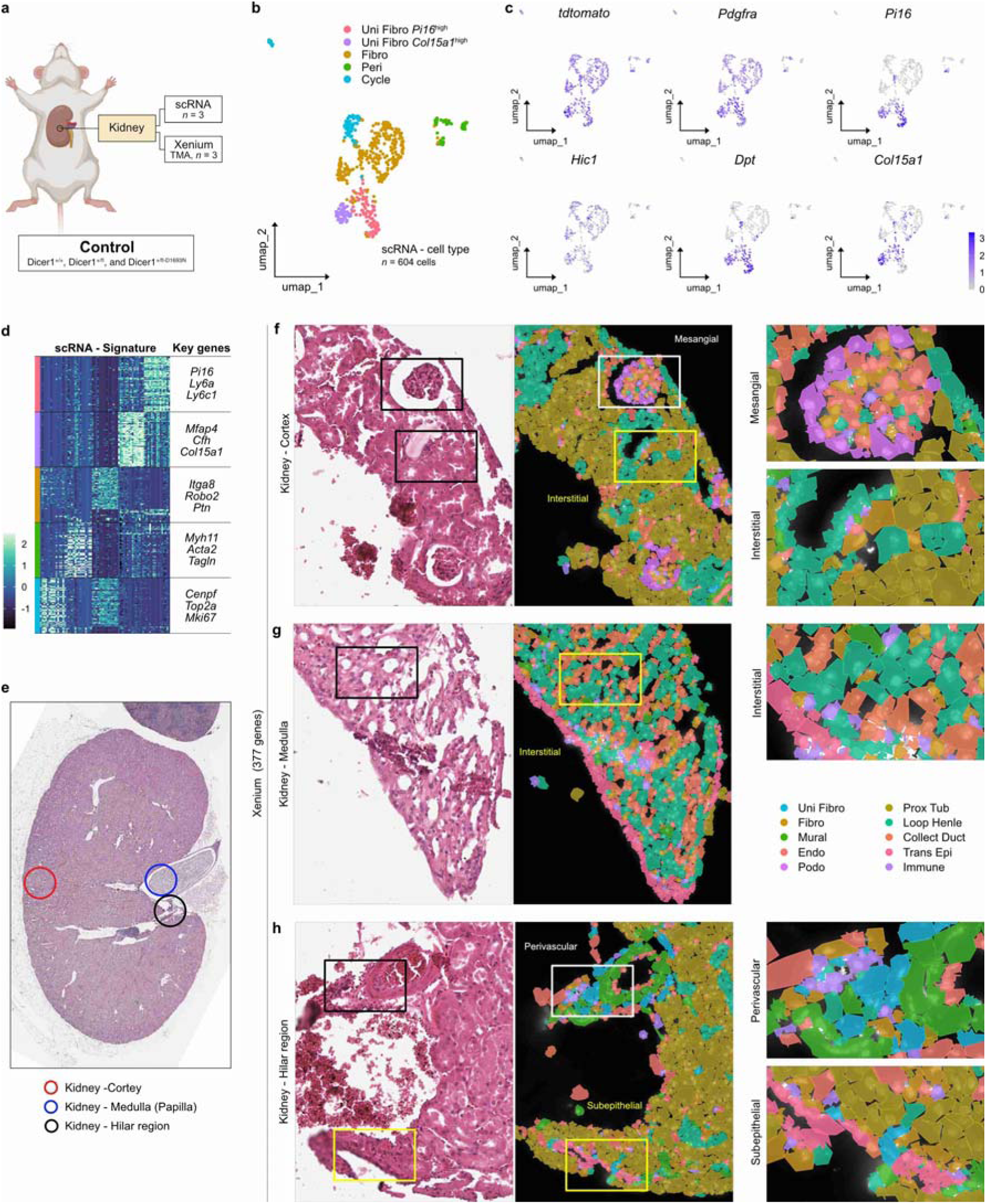
Organization of the Hic1+ mesenchymal stromal cell lineage in murine steady state kidney. **a**, Steady-state kidney samples from control mice (Dicer1^+/+^, Dicer1^+/fl^, and Dicer1^+/fl-D1693N^) profiled by scRNA-seq and Xenium spatial transcriptomics analyses. Created with content from BioRender.com. **b**, UMAP plot from scRNA-seq data (*n*=3) of *tdtomato*^+^ mesenchymal stromal cells colored by cell type. Louvain method was used for clustering, and cell types are annotated based on transcripts representative of different lineages (relating to Extended Data Fig. 3f): Uni Fibro *Pi16*^high^, universal fibroblast *Pi16*^high^; Uni Fibro *Col15a1*^high^, universal fibroblast *Col15a1*^high^; Fibro, Fibroblast/mesangial cell; Mural, mural cells; Cycle, cycling cells. **c**, UMAP plots depicting expression of key fibroblast lineage markers. **d**, Heatmap of top 30 defining transcripts, based on cell type annotation. Key genes of signature are highlighted. **e-h**, Mapping of clusters identified from targeted spatial transcriptomics (10X Genomics, Xenium, 379 genes; *n*=4; related to Extended Data Figs. 4a-e) to spatial domains of murine kidney (**e**) shows presence of Fibro in the mesangium and interstitium of the renal cortex and medulla (**f** and **g**). Aggregation of Uni Fibro is noted in perivascular and basement membrane-associated niches (**h**). Cell types are colored according to the legend.

Spatial analysis of kidney tissue corroborated the scRNA-seq data, revealing a steady-state kidney composition that included MSCs (Extended Data Figs. 4a-e, Supplementary Table 7). Targeted analysis of kidney MSCs (Fig. 2e) identified mural cells (Mural) mapping to pericytes and vascular smooth-muscle cells (*Rgs5^+^*, *Myh11*^+^, *Tagln*^+^), fibroblasts/mesangial cells (Fibro: *Bgn*^+^, *Cfh*^+^, *Itga8*^+^), and universal fibroblasts (Uni Fibro: *Bgn*^+^, *Lum*^+^, *Mfap4*^+^) which appeared to overlap with the Uni Fibro *Pi16*^high^ and Uni Fibro *Col15a1*^high^ populations. In line with their expression profile, Fibro mapped to interstitial fibroblasts of cortex and medulla, as well as to rare mesangial cells (Figs. 2f and g), while Uni Fibro was predominantly found subjacent to transitional cell epithelium and within the perivascular niche (Fig. 2h). Although the resolution of the targeted gene panel did not allow for subclassification of Uni Fibro based on UMAP, *Pi16* expression was notably enriched in Uni Fibro within the perivascular niche (Extended Data Fig. 4f). A recent study of murine fibroblasts also identified two groups of *Dpt*^+^ fibroblasts that are present universally across various tissues (in this paper kidney was not studied)^27^. The first group included *Dpt*^+^, *Pi16*^+^ universal fibroblasts that shared features with adventitial stromal cells^28^, which are found in perivascular niches^29,30^. The second group was composed of *Dpt*^+^, *Col15a1*^+^ specialized fibroblasts and showed an association with the basement membrane. Our study identifies two distinct types of universal fibroblast populations in the steady-state murine kidney, with higher abundance in juvenile mice. We further suggest that *Pi16^high^* universal fibroblasts are typically enriched within the perivascular niche, while *Col15a1^high^*universal fibroblasts are more associated with the subepithelial basement membrane of lining transitional epithelium.

### A Hic1^+^ universal fibroblast lineage as cell of origin for Dicer1 sarcoma

Next, we characterized the transcriptional landscape of primary HDT renal tumor samples by scRNA-seq (*n*=5) (Extended Data Fig. 5, Supplementary Table 8). Unsupervised clustering of *tdTomato*^+^ MSCs (Extended Data Fig. 6a and b) revealed distinct cell populations, including steady-state kidney cell types (Fig. 3b-d, Supplementary Table 9). Analyzing the expression of MSC markers (Extended Data Fig. 6c) and co-clustering of HDT tumor samples and steady-state samples established their molecular similarities (Extended Data Fig. 6e and f). In HDT tumor samples identified populations included two cluster corresponding to steady-state mural cells, including pericytes (Peri; *Rgs5*^+^) and vascular smooth muscle cells (SM Vasc; *Acta2*^+^), as well as a cluster of tumor associated fibroblasts (Fibro Perturb) not identified in steady-state kidney that expressed markers of *Pi16*^high^ universal fibroblasts but were mainly characterized by a distinct gene signature of perturbed-state fibroblasts (*Comp*^+^, *Cpxm2*^+^, and *Itgbl1*^+^)^27^. Additionally, we identified a cluster overlapping with steady-state *Pi16*^high^ and *Col15a1*^high^ universal fibroblasts, that we termed “Progenitor” (Progen), and a “ground” population (Ground) closely resembling Fibro of steady-state kidney. A “proliferative” cluster (Prolif) enriched in genes related to proliferation and cell cycle similar to cycling cells of steady-state kidney was also identified, including cells with both underlying Progen and Ground signatures. Two subpopulations absent in steady-state kidney, expressing genes linked to skeletal muscle development (e.g. *Pax7*) and committed muscle cells (e.g. *MyoG*), termed “transiting-differentiated myogenic cells” (TR Diff Myo) and “differentiated myogenic cells” (Diff Myo), were also found^31^. For trajectory analysis, cells were ordered in pseudotime, with the earliest point set at the node in Progen with expression of *Dpt*^+^ (Extended Data Fig. 6d). Trajectory analysis from the Progen cluster revealed eight trajectories, seven of which led into tumor-associated clusters, accompanied by a loss of Progen-specific markers (e.g. *Dpt*, *Mfap4*) (Fig. 3e). Trajectories 4 led into Diff Myo (*Myog*^+^), passing through the “intermediate state” TR Diff Myo cells (*Pax7*^+^), including a highly proliferative state of TR Diff Myo. Trajectories 1 and 5 transitioned into the Ground state (*Itga8*^+^).

**Fig. 3:**
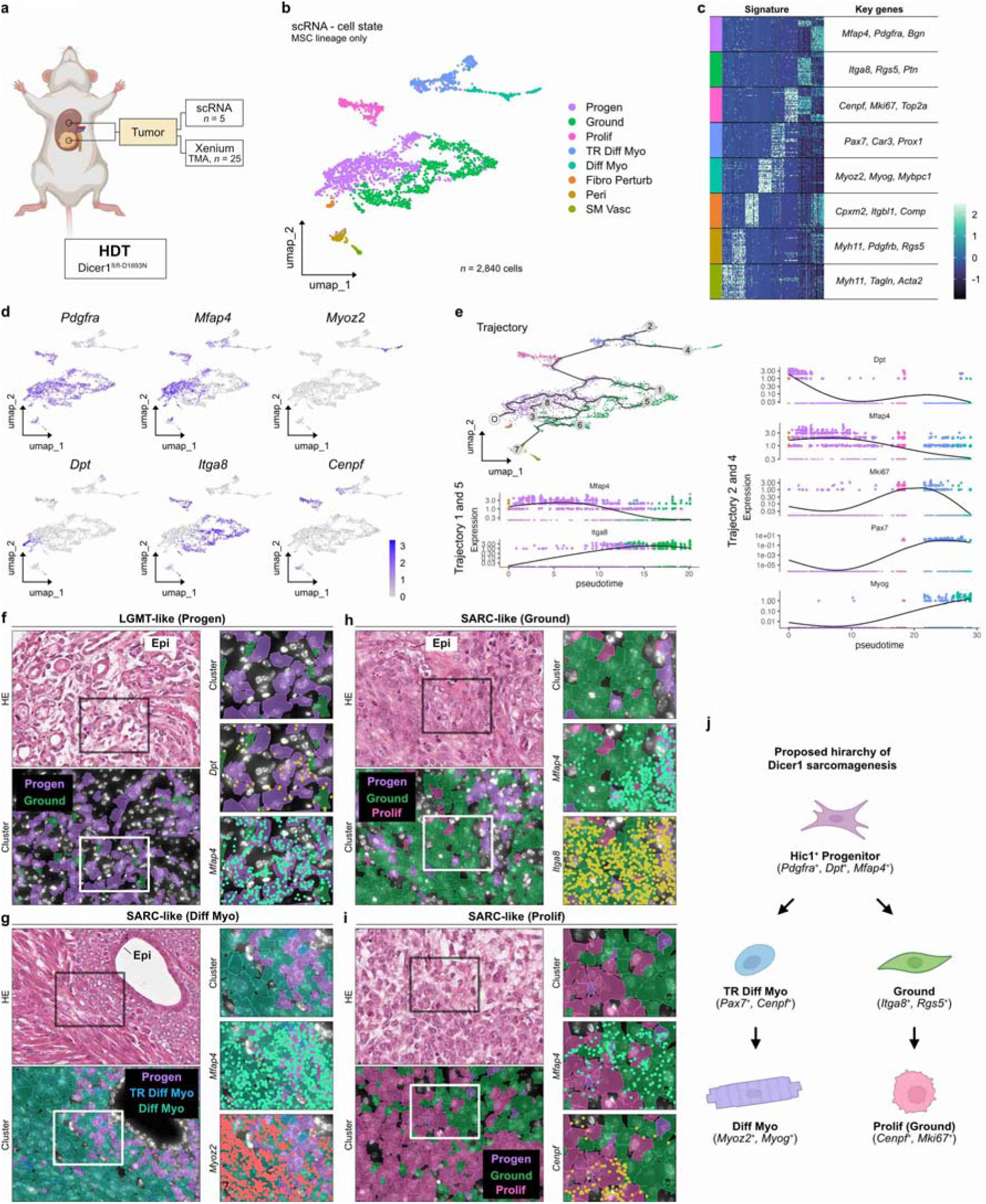
A Hic1+ universal fibroblast lineage as cell of origin for Dicer1 sarcoma. **a**, Samples from HDT mice (Dicer1^fl/fl-D1693N^) profiled by scRNA-seq and Xenium spatial transcriptomics analyses. Created with content from BioRender.com. **b**, UMAP plot from scRNA-seq data (*n*=5) of *tdtomato*^+^ mesenchymal stromal cells colored by cell type. Louvain method was used for clustering, and cell types are annotated based on transcripts representative of different lineages (relating to Extended Data Fig. 5c): Progen, progenitor; Ground, ground (Fibro-like); Prolif, proliferative; TR Diff Myo, transiting-differentiated myogenic cells; Diff Myo, differentiated myogenic cells; Fibro Perturb, perturbed fibroblast; Mural, mural cells. **c**, Heatmap of top 30 signature defining transcripts, based on cell state annotation. Key genes of signatures are highlighted. **d**, UMAP plots depicting expression of key markers. **e**, Trajectory analysis of scRNA-seq data. UMAP colored by cell type. O indicates origin, trajectories are numbered. Gene trend plots of key cluster genes along the sarcomagenic trajectory (Ground and Diff Myo) defining the transformation process. Cells are colored according to cell type. **f-i**, Mapping of clusters identified from targeted spatial transcriptomics (10X Genomics, Xenium, 379 genes; *n*=25; related to Extended Data Figs. 7) to spatial domains of HDT tumors. Progen is abundant in tumors with LGMT-like morphology, shows primitive cytologic features and expresses *Dpt* and *Mfap4* (**f**). In SARC-like tumors with rhabdomyoblastic differentiation, Progen colocalizes with TR Diff Myo and Diff Myo. Expression of *Mfap4* and *Myoz2* is shown (**g**). Ground corresponds to sarcomatous cells, which may be colocalized with Progen (**h**) and may progress into highly atypical/anaplastic and proliferative tumor cell populations (Prolif). Expression of *Mfap4*, *Itga8*, and Cenpf is shown (**i**). **j**, Schematic of proposed hierarchy of Dicer1 sarcomagenesis. Created with content from BioRender.com.

To spatially correlate these clusters, we performed targeted spatial transcriptomics (10X Genomics, Xenium, 377 genes) on 25 HDT samples, revealing clusters corresponding to both steady-state kidney and tumor-associated cell states, consistent with our scRNA-seq data (Extended Data Fig. 7, Supplementary Table 10). Spatial analysis revealed Progen cluster cells in samples with LGMT-like morphology, expressing *Dpt* and *Mfap4* (Fig. 3f), supporting their critical role during cancer initiation. Moreover, we demonstrated a close spatial relationship between the cell state trajectories observed in scRNA-seq data, thereby validating their biological relevance and confirming the consistency of these trajectories with underlying tissue architecture (Figs. 3g and h). Notably, many cells assigned to Ground, and especially Prolif clusters showed frank sarcomatous differentiation (Fig. 3i).

Based on these findings, we propose a developmental hierarchy for Dicer1 sarcomagenesis, beginning with a Hic1^+^ MSC that shares features with *Pdgfra*^+^, *Dpt*^+^*, Mfap4*^+^ universal fibroblasts (Fig. 3j). This progenitor population gives rise to a myogenic lineage, progressing through a proliferative *Pax7*^+^ transitional state before differentiating into committed myogenic cells. Alternatively, the progenitor population can give rise to a sarcomatous ground state, transcriptionally resembling non-universal fibroblasts/mesangial cells of steady-state kidney, as well as a highly proliferative cell state, both associated with frank sarcomatous differentiation.

### Murine cell states are largely recapitulated in human DICER1 sarcoma

Human DICER1-associated mesenchymal neoplasms closely mimic RMS clinically and morphologically. However, our previous molecular analysis distinguished DICER1 sarcomas from bona-fide RMS subtypes (fusion-negative, fusion-positive and *MYOD1*-mutated RMS) based on DNA methylation profiles, suggesting distinct cellular origins for these tumors^9^. To build on these findings, we characterize the transcriptional landscape across human DICER1 mesenchymal tumors (LGMT DICER1, SARC DICER1, and PIS DICER1), including Cystic nephroma, pleuropulmonary blastoma (PPB) I/II/III, nasal chondroid mesenchymal hamartoma, DICER1-mutated embryonal RMS, and primary intracranial sarcoma (PIS DICER1) (Fig. 4a). We analyzed 16 patient samples using a targeted spatial transcriptomics platform (10X Genomics, Xenium, 377 genes), performing unsupervised clustering that identified 13 Louvain clusters (Extended Data Figs. 8a and b). These clusters correspond to distinct cell populations, including non-neoplastic cell types (Extended Data Figs. 8c and d). Guided by lineage markers (e.g. *PDGFRA*) and spatial morpho-molecular correlation, we identified fibroblast lineage clusters (Extended Data Fig. 8e)^32^, which were grouped into cell states based on shared transcriptomic characteristics, largely recapitulating cell states identified in our murine HDT tumors (Figs. 4b-d, Supplementary Table 11). These included a neoplastic “progenitor” (Progen) cluster expressing *PDGFRA* and human universal fibroblastic marker *DPT*, a “proliferative” (Prolif) cluster dominated by proliferation and cell cycle genes, two myogenic subpopulations expressing skeletal muscle-related genes (e.g. *DES*, *PROX1*), as well as committed muscle cells (e.g. *MYBPC1*), termed “transiting-differentiated myogenic cells” (TR Diff Myo) and “differentiated myogenic cells” (Diff Myo). Additionally, a *PDGFRA*^+^ fibroblastic “ground” population lacking other signatures, and a distinct fibroblast cluster (Fibro) mapping to non-neoplastic tissue and marked by *PDGFRA*^+^ and *C7*^+^ were identified. Spatial analysis revealed Progen cells in all tumors at varying frequencies (Fig. 4e), often mapping to the cambium layer of LGMT and SARC DICER1 (Figs. 4f). Like HDT tumors, TR Diff Myo states were associated with increased proliferation (Fig. 4g). Bona fide rhabdomyoblastic differentiation (Diff Myo) co-localized with TR Diff Myo and the committed muscle marker *MYBPC1* (Fig. 4h). Ground and Prolif cells were abundant in SARC/PIS DICER1 and exhibited sarcomatous growth (Fig. 4i). These results alongside trajectory analysis (Extended Data Fig. 8f) closely mirrored spatial relationships observed in murine HDT tumors.

**Fig. 4:**
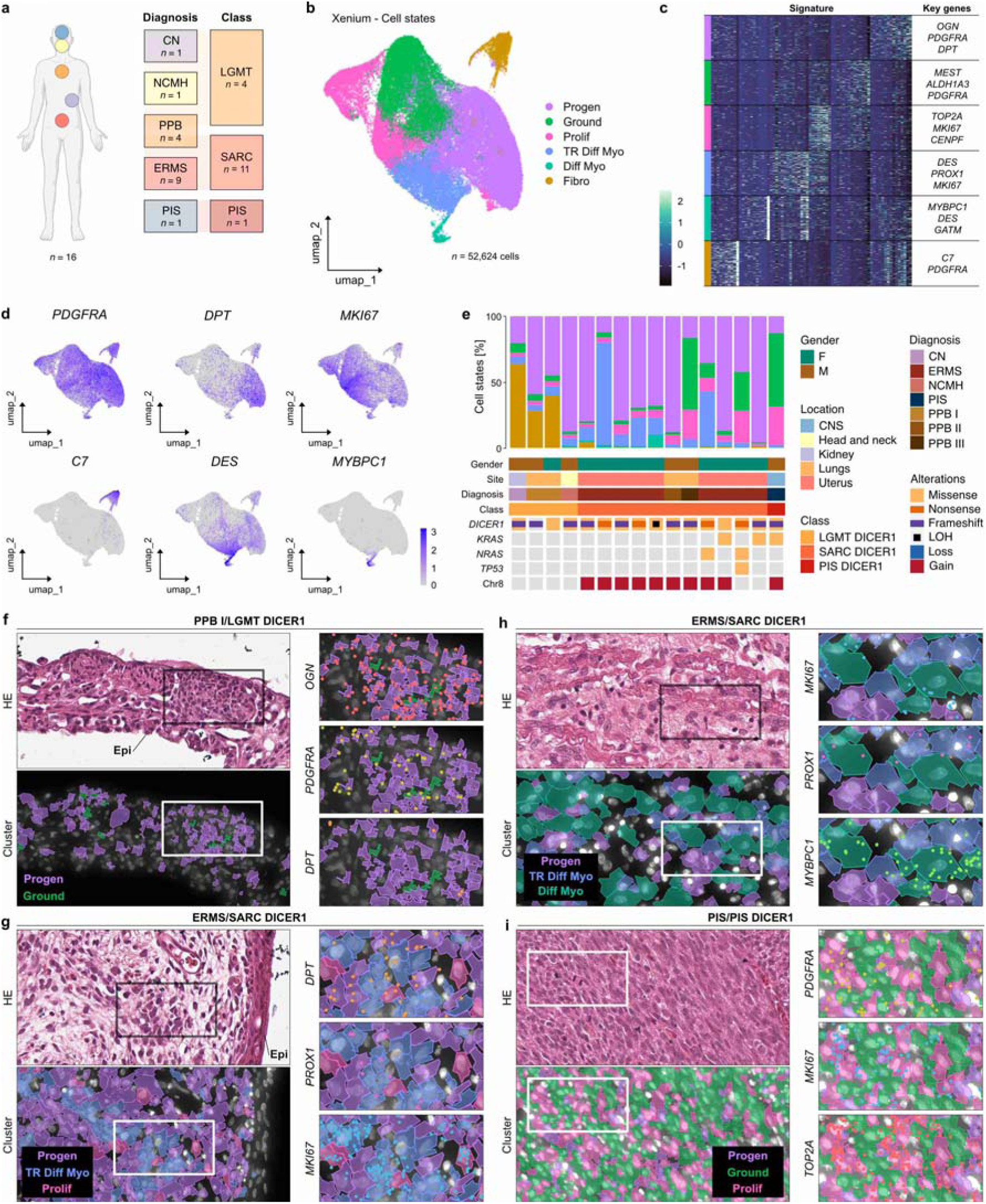
Murine cell states are largely recapitulated in human DICER1 sarcoma. **a**, Patient samples (*n*=16) encompassing well established tumor entities comprising three molecular classes of DICER1-associated mesenchymal neoplasms^9^. Created with content from BioRender.com. **b,** UMAP plot from spatial transcriptomic data of mesenchymal cells from fibroblastic lineage colored by cell type. Louvain method was used for clustering, and cell types are annotated based on transcripts representative of lineage (relating to Extended Data Fig. 8d) and morpho-molecular correlation (relating to panels f-i): Progen, progenitor; Ground, ground; Prolif, proliferative; TR Diff Myo, transiting-differentiated myogenic cells; Diff Myo, differentiated myogenic cells; Fibro, fibroblasts. **c**, Heatmap of top 15 signature defining transcripts, based on cell state annotation. Key genes of signatures are highlighted. **d**, UMAP plots depicting expression of key fibroblast lineage markers and markers characteristic of identified cell types. **e**, Summary of the cell type composition of each patient sample with clinical and tumor sequencing information included as an oncoplot below. Class: low-grade mesenchymal tumor with DICER1 alteration, LGMT DICER1; sarcoma with DICER1 alteration, SARC DICER1; primary intracranial sarcoma with DICER1 alteration, PIS DICER1. Diagnosis: cystic nephroma, CN; embryonal rhabdomyosarcoma (DICER1-mutated), ERMS; nasal chondroid mesenchymal hamartoma, NCMH; primary intracranial sarcoma, PIS; pleuropulmonary blastoma type I, II, and III, PPB I/II/III. LOH, loss of heterozygosity. **f-i**, Mapping of clusters identified from targeted spatial transcriptomics to spatial domains of human tumors. PPB I (LGMT DICER1) shows a cystic configuration with a septum and nodular and subepithelial growth of primitive tumors cells which are assigned to Progen. Expression of *OGN*, *PDGFRA*, and *DPT* is shown (**f**). Example of a DICER1-mutated ERMS (SARC DICER1) with a subepithelial cambium layer and focal aggregation of more atypical tumor cells that map to TR Diff Myo. Expression of *DPT*, *PROX1*, and *MKI67* is shown (**g**). Example of another DICER1-mutated ERMS (SARC DICER1) with bona fide rhabdomyoblastic differentiation and colocalization of cells assigned to TR Diff Myo and Diff Myo. Expression of *MKI67*, *PROX1*, and *MYBPC1* is shown (**h**). PIS with fascicular and frank sarcomatous growth of atypical and highly proliferative tumor cells assigned to Ground and Prolif clusters. Expression of *PDGFRA*, *MKI67*, and *TOP2A* is shown (**i**).

A recent scRNA-based meta-analysis identified a robust tripartite cell state landscape across various human RMS tumors, including “Progenitor”, “Proliferative”, and “Differentiated” metaprograms^33^. This paper also inspired some of our cluster names. Our transcriptomics data showed that these RMS metaprograms are enriched in signatures present in DICER1 mesenchymal neoplasms (Extended Data Fig. 8g), highlighting their transcriptional relatedness.

Although our analysis is limited by the targeted panel, the striking similarity between cell states identified in murine and human DICER1 tumors further substantiates the relevance of our GEMM model of DICER1 sarcoma. This reinforces its fidelity to the human disease and underscores its potential as a tool for advancing mechanistic understanding and developing novel therapeutic strategies.

## Discussion

Patients with DICER1 syndrome are predisposed to a variety of cancers, many of which are aggressive sarcomas that arise in diverse anatomical locations. Recent multi-omic studies have significantly advanced our understanding of the genetic underpinnings of these cancers. However, the specific factors driving tumor initiation remain poorly understood. In this study, we demonstrate that expression of a compound heterozygous *Dicer1* mutation in Hic1^+^ MSC is associated with the development of renal sarcomas that closely resemble the human disease, both histomorphologically and genomically. These tumors recapitulate key secondary genetic events involving *Kras* and *Trp53*, mirroring alterations observed in human DICER1-associated sarcomas, including anaplastic sarcoma of the kidney^9,18–21^.

Our findings suggest that Dicer1 disruption in Hic1^+^ MSCs has a high potential for the development of Dicer1-associated sarcoma. By using a lineage-traceable GEMM, we identified Hic1^+^ MSCs marked by *Pdgfra*, *Dpt*, and *Mfap4* as putative cells of origin. In steady-state murine kidneys, we show that *Dpt*^+^ universal fibroblasts are localized to the perivascular and subepithelial basal membrane niche. This is particularly intriguing given that DICER1 sarcomas frequently arise in organs with epithelial lining, such as the lungs, kidneys, and uterus, and form a cambium layer (sarcoma botryoides)^2^. Our data suggest that the anatomical predilection for DICER1 sarcomas for subepithelial regions may be linked to the proposed cellular origin within the universal fibroblastic niche associated with the basal membrane. However, the restriction of tumor development to the kidney in our murine model remains unclear. While universal fibroblasts exist across a variety of tissues in mice^27^, we hypothesize that the specific cell states of universal fibroblasts in the kidney may influence the susceptibility to Dicer1-associated transformation. Indeed, our analysis of MSCs in steady-state kidney revealed that the universal fibroblast populations vulnerable to transformation are more abundant in juvenile mice compared to adults. Given that tumors from the DICER1 sarcoma spectrum predominantly affect children and young adults, we further hypothesize that tumor susceptibility in humans may also be influenced by developmental stage, with earlier developmental windows rendering certain fibroblast populations more prone to malignant transformation.

We further demonstrate that the epithelial lining in our kidney tumors was negative for tdTomato, indicating that it did not belong to the neoplastic lineage. Nevertheless, these epithelial areas frequently displayed a botryoid or even cystic appearance. We propose that such cystic changes, commonly observed in tumors from the DICER1 mesenchymal neoplasm spectrum - such as PPB I and CN - are not a result of DICER1-driven neoplastic epithelial growth, as previously suggested^4,13^. Instead, we hypothesize that these cystic transformations are likely a reactive phenomenon, driven by neoplastic mesenchymal proliferation, which may exert pressure on adjacent structures, such as kidney tubules or alveolar spaces, leading to their compression and subsequent cystic dilation due to fluid or air entrapment.

Upon induction of a compound heterozygous *Dicer1* mutation, we observed the expansion of the fibroblastic progenitor population that gave rise to divergent trajectories, including a step-wise continuum of rhabdomyoblastic differentiation. Notably, our HDT tumors closely mimic conventional types of RMS morphologically and transcriptionally. Conventional RMS subtypes are typically thought to originate from satellite-cell derived post-natal muscle and results from aberrant skeletal muscle differentiation^34,35^. However, recent studies have challenged this notion, suggesting that RMS may originate from endothelial progenitors^36^ or mesenchymal stem cells^33,37^. Interestingly, a recent study demonstrated that fibroblasts derived from the lateral plate mesoderm can trans-differentiate by activating myogenic programs during physiological development^38^. Yaseen et al. showed that these fibroblasts not only acquire myogenic features and fuse with existing myofibers but also retain partial expression of their original fibroblastic transcriptional program. In line with this, our findings provide clear evidence that fibroblast have the potential to undergo rhabdomyoblastic transformation upon *Dicer1* disruption, challenging the long-standing dogma that RMS tumors develop from a predetermined myogenic lineage.

Finally, our data reveals a striking concordance between the cellular origins of murine and human DICER1 sarcomas. We confirm that in the human disease, cancer cell states share transcriptional similarities with those observed in our murine model, including the presence of a progenitor population of universal fibroblastic lineage (*PDGFRA*^+^, *DPT*^+^). Additionally, we demonstrate that DICER1-associated mesenchymal neoplasms exhibit transcriptional similarities to other tumors within the RMS spectrum, highlighting their relatedness.

Overall, our study provides a deeper understanding of the cellular contexts driving DICER1-associated sarcomagenesis and opens promising avenues for future mechanistic investigations and therapeutic development for tumors of the RMS spectrum.

## Authors’ Disclosures

The authors declare no conflicts of interest.

## Authors’ contributions

F.K.F.K.: resource, methodology, investigation, formal analysis, visualization, writing; J.Y.Z.: methodology, investigation, formal analysis, writing; S.C. and J.S.: Methodology, investigation; B.L.: formal analysis, data curation, visualization; Y.M., L.A.H. and J. B.: investigation; W.S.: methodology; K.S.C.: discussion and manuscript editing; A.v.D: resource; W.F.: resource; T.M.U.: resource, supervision, manuscript editing; Y.W.: conceptualization, methodology, investigation, formal analysis, writing, supervision, project management, funding acquisition. D.G.H.: conceptualization, writing, supervision, funding acquisition.

## Acknowledgements

We appreciate the technical support of Chae Young Shin, Raymond Zirui Feng, Dionzie Ong, Rebecca Ho, Eunice Li, Emma Guo, Christine Chow, Yimei Qin and all staff in the Animal Research Centre of British Columbia Cancer Research Institute. We would also like to express our gratitude to the collaborating clinicians, as well as to the patients who generously provided their samples for this research. This project was supported by research funds from a Terry Fox New Frontiers Program Project Grant (#1082, DGH), Canadian Institutes of Health Research (CIHR) grants (MOP-154290, D.G.H.; PJT#178179, Y.W.; PJT#195754, D.G.H. and Y.W.), and the philanthropic supports to British Columbia Ovarian Cancer Research Centre through the BC Cancer Foundation and the VGH/UBC foundation. F.K.F.K. is funded by the Deutsche Forschungsgemeinschaft (DFG, German Research Foundation) – Projektnummer 523898075. J.Y.Z. is a recipient of CIHR CGSD. D.G.H. is supported by a Canadian Research Chair in Molecular and Genomic Pathology.

## Methods

### Model development and cohort assembly

All animal experiments were conducted following ethical approval by the University of British Columbia animal care committee (A17-0054, A17-0055, A21-0023 and A21-0047). The conditional knockin mice expressing the floxed Dicer1 p.D1693N (Dicer1^+/fl-D1693N^) was previously developed by our group^10^. The Dicer1^fl/fl^ mice^33^ [RRID:MMRRC_015388-UNC] were obtained from the Mutant Mouse Resource & Research Centres (MMRRC) depository. The (Rosa26)^LSL-tdtomato^ strain was obtained from The Jackson Laboratory (stock #: 007909). Hic1^CreERT2^ mice were previously generated as described elsewhere^15^. Mouse lines were maintained on a pure C57Bl/6J genetic background. We outcrossed these 4 strains together to develop the Hic1^CreERT2^;Dicer1^fl/fl-D1693N^;(Rosa26)^LSL-tdTomato^ (HDT) strain. Genotyping was performed using primers in Supplementary Table 12.

To induce *Dicer1* mutations, mice were injected intraperitoneally with tamoxifen (dose: 1mg/kg 5 consecutive days for mice > 3 weeks old; 2mg/kg 3 consecutive days for mice <2 weeks old). The cohort included wild type control mice and mutant HDT mice that received a tamoxifen injection within first 2 months of birth (<=61 days). Mice were either found dead or sacrificed because they were moribund (defined as grade 4, 5 sick reports by BC Cancer Research Centre ARC facility) or reached the experimental end point (1.5 years). A subset of non-moribund HDT mice were sacrificed for interim assessment. A subset of control mice was sacrificed for scRNA-seq time point analyses. For penetrance analysis, the presence of macroscopic or microscopic tumors at death was used as the primary endpoint. For overall survival (OS) analysis, events were defined as either the death of the mice or their euthanasia due to moribund conditions.

### Patient samples

Patient samples of DICER1 mesenchymal neoplasms included in this study (CN, embryonal RMS, NCMH, PIS DICER1, and PPB) had all undergone histopathological evaluation by specialized pathologists, as well as molecular testing whenever applicable. Samples were collected at McGill University-affiliated hospitals in Montreal, Quebec, Canada and the Institute of Pathology, University Hospital Heidelberg, mainly from collaborating institutions in accordance with ethics review board regulations. Eligible participants signed an informed consent form. Clinical data, histological features and/or sequencing results of tumors have previously been reported elsewhere^9^. Study of patient samples was approved by the institutional ethics committee (Heidelberg University, S660/2020) and performed in accordance with the Declaration of Helsinki.

### Tissue microarray construction

For murine tissue microarray (TMA) construction, 1.0_mm cores of formalin-fixed, paraffin-embedded (FFPE) tissue from control and HDT mice were used, as described previously^39^. For human samples, a previously constructed TMA was used, that included duplicates of 0.6 mm cores of FFPE tissues, including the 16 samples of DICER1 mesenchymal neoplasms used in this study.

### Histo- and Immunohistochemistry

4 µm-sections were cut and mounted on Permaflex Plus Microscopic Slides (Leica Biosystems). Sections were stained using standard protocols for Hematoxylin and Eosin (H&E). For immunohistochemistry of murine samples slides were subjected to heat-induced antigen retrieval in a high-pH buffer. Staining was performed using the Ventana Benchmark Ultra automated immunostainer (Ventana Medical Systems, Tucson, AZ, USA). Immunohistochemical staining of murine samples was performed with antibodies to tdTomato (Rockland, anti-RFP, 600-401-379, Dilution 1:400), Pdgfra (Cell signaling, D1E1E, Dilution 1:100), and Desmin (Abcam, Y66, ready-to-use).

### Targeted panel sequencing

Genomic DNA was extracted with QIAamp DNA FFPE Advanced UNG Kit (Cat. # 56704). Mutations in *Kras* (exons 2 and 3), *Nras* (exons 2 and 3), *Hras* (exons 2 and 3), *Trp53* (exons 2-11), and *Dicer1* (exon 24) were sequenced on a NovaSeq 6000 machine using in house designed primers, and libraries were prepped with in house. Targeted panel data was processed using a modified version of the Nextflow^40^ and Sarek^41^ pipeline, version 3.4.2. Raw reads were trimmed with FastP and aligned with bwa-mem2. Overlapping primer intervals were merged with bedtools merge, version 2.30.0, and 100 bp of padding was added to either side of each interval. No duplicate marking was carried out and the ‘wes’ parameter was set to true. Somatic variants were called on the primer intervals with GATK Mutect2 with ‘ignore_soft_clipped_bases’ set to true, and annotated with SnpEff^42^ and Ensembl VEP^43^. For the variant calling step, the pipeline was modified to disable downsampling of the reads. Variants of interest were selected by filtering for those with variant allele fraction (VAF) > 0.1, and excluding those annotated as ‘synonymous_variant’ or ‘intron_variant’ by SnpEff.

### *Kras* qPCR and FISH

To evaluate *Kras* copy number status we used a qPCR assay with SYBR Green detection and a custom primer for *Kras* (Chr.6:145220129) and *Tfrc* (Chr.16:32626732). Samples were normalized to *Tfrc* and evaluated for a relative gain in *Kras*. HDT361 showed a high count for *Kras* upon qPCR, and gene amplification was identified using a fluorescence in-situ hybridization (FISH) assay. For this assay we designed custom probes for *Kras* on 6qA1 (RP23-300B22, green; Empire genomics) and 6qG3 (RP23-118J6, red; Empire genomics).

### scRNA-seq

#### Sample preparation

Tumour and normal kidney tissues were minced to 1mm^3^ with forceps and incubated at 37°C with Stemcell 10X Collagenase/hyaluronidase in DMEM (10X Genomics, C#07912) in HBSS for 30 minutes. The minced tissues were subsequently incubated in Trypsin-EDTA (0.25%) for 5 min at 37°C. Cells were then filtered through a 70 μm cell strainer. Live cells were enriched with Miltenyi Biotec Dead Cell Removal Kit. Mesenchymal cell populations were then isolated by FACS sorting with tdTomato positive enrichment and antibody depletion of Epcam, CD45, CD31, CD11b and F4/80. Cells were then prepared according to the 10X Genomics Chromium Single Cell Universal 3‘ Gene Expression kit protocol.

#### Alignment and quality control

Raw reads were aligned with Cell Ranger count 7.1.0 to a combined reference genome, composed of the mm10 reference with the addition of the tdTomato reporter sequence (REF Underhill Hic1). The combined reference genome was generated with Cell Ranger mkref 7.1.0. Following alignment, per-sample quality control was conducted. Low quality cells with > 25% of reads originating from mitochondrial genes, < 1000 UMIs detected, or < 500 genes detected, were discarded.

#### Integration, clustering, cell-type annotation, and pathway activity

Samples originating from HDT or control mice were integrated with other samples of their respective genotype using Seurat 4.3.0.1^44^. Counts were normalized per cell by library size followed by log1p transformation. The top 2,000 highly variable genes per sample were selected using the VST method and the top 2,000 genes across samples were used for integration via reciprocal principal components analysis (PCA). After integration and dimensionality reduction via PCA, shared nearest neighbors were identified in PCA space and clustered with the Louvain algorithm, using the first 30 components. The resulting clusters were manually annotated via the examination of differentially expressed up-regulated genes, in combination with markers of kidney and Hic1^+^ MSC types obtained from the literature. To achieve a pure Hic1^+^ MSC lineage following examination of all cell types, the data was subset to the relevant clusters in the control (Uni Fibro, Fibro, Mural, Cycle) or HDT (Fibro Perturb, Fibro, Mural, Cycle, TR Diff Myo, Diff Myo) samples. If necessary, residual *tdTomato*-cells, or cells expressing epithelial (*Epcam*^+^, *n*=102), immune (*Ptprc^+^*, *n*=113), or transitional epithelial (*Krt19*^+^, *n*=376) markers were excluded from the subset data. The resulting data, filtered for the Hic1^+^ MSC lineage, was reprocessed as described above for the unfiltered data; a reduced number of principal components were used for neighbor finding and clustering after selection via an elbow plot.

#### Trajectory inference

The Hic1^+^ MSC lineage obtained from the HDT samples was used as input for trajectory inference with Monocle 3, version 1.3.4^45^. Based on the UMAP obtained during dimensionality reduction with Seurat, trajectory was inferred, and the cells were placed in a pseudotime ordering, setting the node in the graph surrounded by the greatest number of *Dpt*^+^ cells as the earliest point. Trajectories to Ground, Cycle, and Diff Myo were examined by selecting the relevant branches, holding the start point constant and varying the end point to the most distant node in the ending cluster of interest. Selected genes of interest were visualized in pseudotime over the respective trajectories.

### Spatial transcriptomics (Xenium)

#### Slide Preparation and Imaging

For spatial transcriptomic analyses (10X Genomics Xenium), two slides were prepared. One slide included a section of the murine TMA, and a second slide featured a section of the human TMA. Tissue sections were prepared as per the Xenium In Situ for FFPE – Tissue Preparation Guide (10X Genomics, CG000578 version Rev C) protocol. 5 μm sections were cut and mounted onto the Sample Area of a Xenium slide. The slides were air dried, followed by a 3 h incubation at 42°C. Subsequently, the slides were processed, over a two-day span, as per the Xenium In Situ for FFPE – Deparaffinization and Decrosslinking (10X Genomics, CG000580 version Rev C) and Xenium In Situ Gene Expression (10X Genomics, CG000582 version Rev C) protocols. On day one, slides were incubated at 60°C for 2 h on a Xenium thermocycler adaptor plate (BioRad, C1000 Touch Thermocycler), followed by deparaffinized through a series of immersions in xylene, 100%, 96% and 70% ethanol, and rehydration with nuclease-free water. Slides were assembled into Xenium cassettes for all downstream steps, starting with decrosslinking and permeabilization. Gene panel probes were hybridized overnight (minimum 18 h) at 50°C. The pre-designed dataset for the murine slide included 379 genes (10X Genomics, Mouse Tissue Atlassing Panel, PN1000627). The pre-designed dataset for the human slide included 377 genes (10X Genomics, Human Multi-Tissue and Cancer Panel, PN1000626). On day two, slides were subjected to a post-hybridization wash (37°C for 2 h), ligation (37°C for 2 h), amplification (30°C for 2 h), autofluorescence quenching and nuclei staining steps. Slides were then stored in PBS-T, overnight, at 4°C, in the dark. The following day, slides and consumables were loaded onto the Xenium Analyzer instrument according to the Xenium Analyzer (10X Genomics, CG000584, version Rev B) protocol. The whole slide area was selected for probe decoding and image acquisition, and the run was initiated. After run completion, slides were H&E stained as per the “Xenium in Situ Gene Expression – Post Xenium Analyzer H&E Staining” (10X Genomics, CG000613, version Rev A) protocol, and scanned using a slide scanner (AperioAT2, Leica).

#### Segmentation and Xenium data analysis

The Xenium bundles were segmented with Xenium Ranger 1.7.1.1, with the expansion distance around nuclei set to 5 µm. The bundles were then re-segmented with Proseg 1.1.4 using the ‘xenium’ preset, converted to the Baysor format with the ‘proseg-to-baysor’ utility bundled with Proseg, and finally re-imported with Xenium Ranger 1.7.1.1^46^. Cell visualization and morpho-molecular correlation was performed in the 10X Xenium Explorer (Version 3.0.1). Resulting Xenium objects were analyzed using Seurat 4.3.0.1^44^. Quality control was conducted and low quality cells with less than 3 counts and less than 3 features, were discarded. After normalization, integration and dimensionality reduction via PCA, shared nearest neighbors were identified in PCA space and clustered with the Louvain algorithm. The resulting clusters were manually annotated via the examination of differentially expressed up-regulated genes, in combination with markers of kidney and Hic1^+^ MSC types obtained from the literature, as well as morpho-molecular correlation. To achieve a pure tumor cell-assoiciated lineage following examination of all cell types, the data was subset to the relevant clusters in the patient samples (Fibro, Progen, Ground, TR Diff Myo, Diff Myo, and Prolif). The resulting data was reprocessed as described above for the unfiltered data; a reduced number of principal components were used for neighbor finding and clustering after selection via an elbow plot. Trajectory inference of patient samples was performed with Monocle 3, version 1.3.4^45^. On the basis of the UMAP obtained during dimensionality reduction with Seurat, trajectory was inferred, and the cells were placed in a pseudotime ordering, setting the node in the graph surrounded by the greatest number of *DPT*^+^ cells as the earliest point.

For comparison of human cell states to metaprograms of RMS tumors^33^, RMS gene signatures were intersected with genes present on the targeted panel. Based on a limited representation of the RMS signatures, the *AddModuleScore* function of Seurat was used to infer metaprogram scores onto human cell states.

## Supplementary Tables Legend

**Supplementary Table 1**: Characteristics of control and HDT mice (*n*=130).

**Supplementary Table 2**: Characteristics of analyzed HDT tumor samples (*n*=57).

**Supplementary Table 3**: Characteristics of scRNA-seq cohort (*n*=12).

**Supplementary Table 4**: Characteristics of Xenium spatial transcriptomics cohort (*n*=29 samples).

**Supplementary Table 5**: DE for cell types identified from unfiltered scRNA of steady-state murine kidney (relating to Extended Data Fig. 3c).

**Supplementary Table 6**: DE for cell types identified from scRNA of steady-state murine kidney limited to MSC lineage (relating to Fig. 2b).

**Supplementary Table 7**: DE for cell types identified from targeted Xenium analysis of steady-state murine kidney (relating to Extended Data Fig. 4c).

**Supplementary Table 8**: DE for cell types identified from unfiltered scRNA of HDT tumors (relating to Extended Data Fig. 5d).

**Supplementary Table 9**: DE for cell types identified from scRNA of HDT tumors limited to MSC lineage (relating to Fig. 3b).

**Supplementary Table 10**: DE for cell types identified from targeted Xenium analysis of HDT tumors (relating to Extended Data Fig. 7e).

**Supplementary Table 11**: DE for cell types identified from targeted Xenium analysis of patient samples limited to fibroblastic populations (relating to Fig. 4b).

**Supplementary Table 12**: Primer sequences for HDT strain genotyping.

## Notes

### Competing Interest Statement

The authors have declared no competing interest.

