## Extended Data Figures for "Spatial single cell transcriptomic analysis of a novel DICER1 Syndrome GEMM informs the cellular origin and developmental hierarchy of associated sarcomas"

**Extended Data Fig. 1**

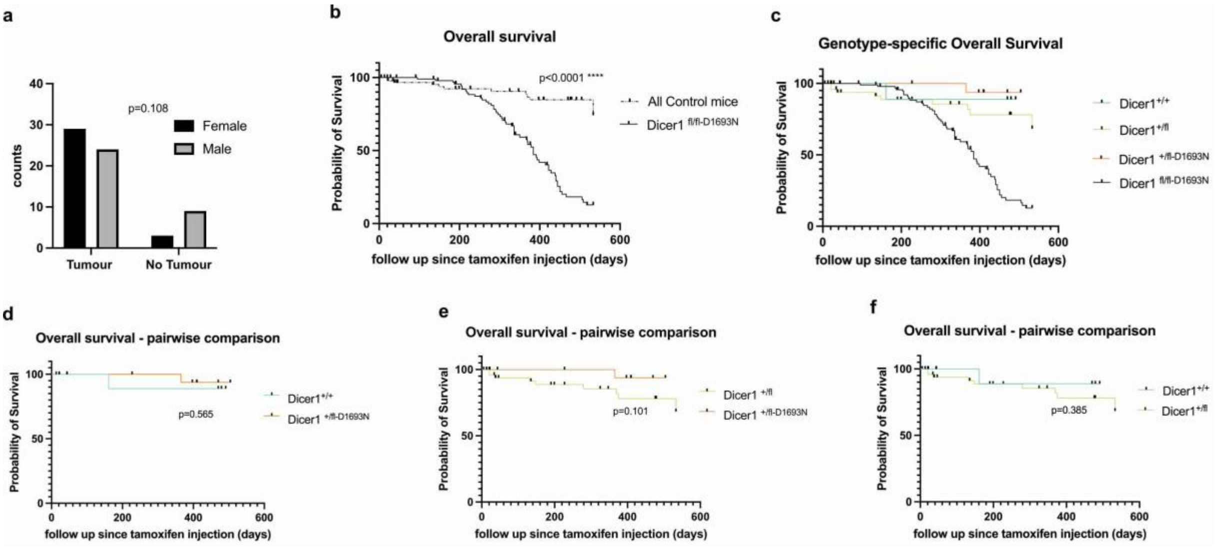


**Extended Data Fig. 1: a**, Comparison of tumor development in HDT mice, stratified by gender using Fisher's exact test, 2 sided (*p*=0.108). **b-f**, Kaplan-Meier estimates for overall survival (OS) of the murine study cohort (referring to Supplementary Table 1), stratified by (**b**) control versus HDT groups, (**c**) genotypes, and pairwise comparisons of OS (**d-f**).

**Extended Data Fig. 2**

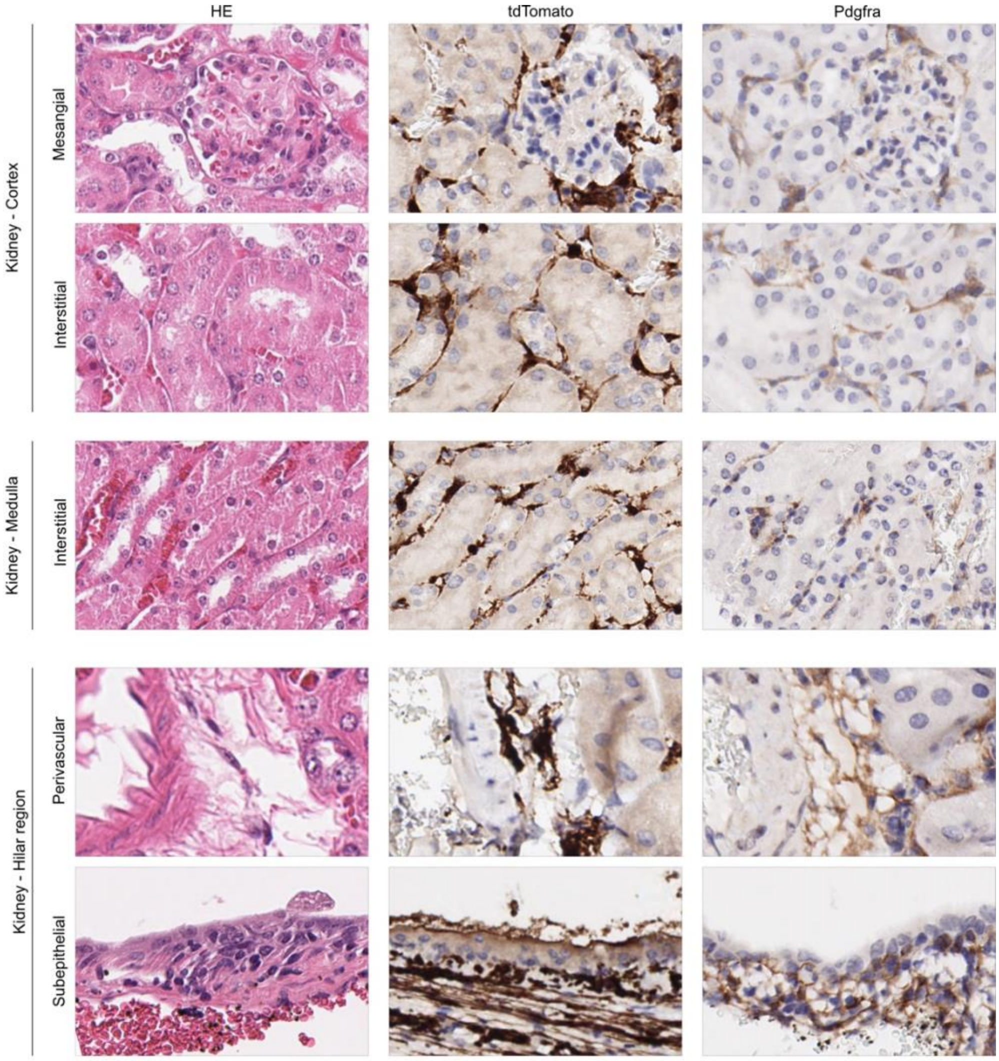


**Extended Data Fig. 2:** Immunohistochemistry for tdTomato and Pdgfra highlight the distribution of MSCs of the Hic1⁺ lineage in the renal cortex, medulla and hilar region, including perivascular and basal membrane-associated niches (relating to Fig. 2e-h).

### Extended Data Fig. 3

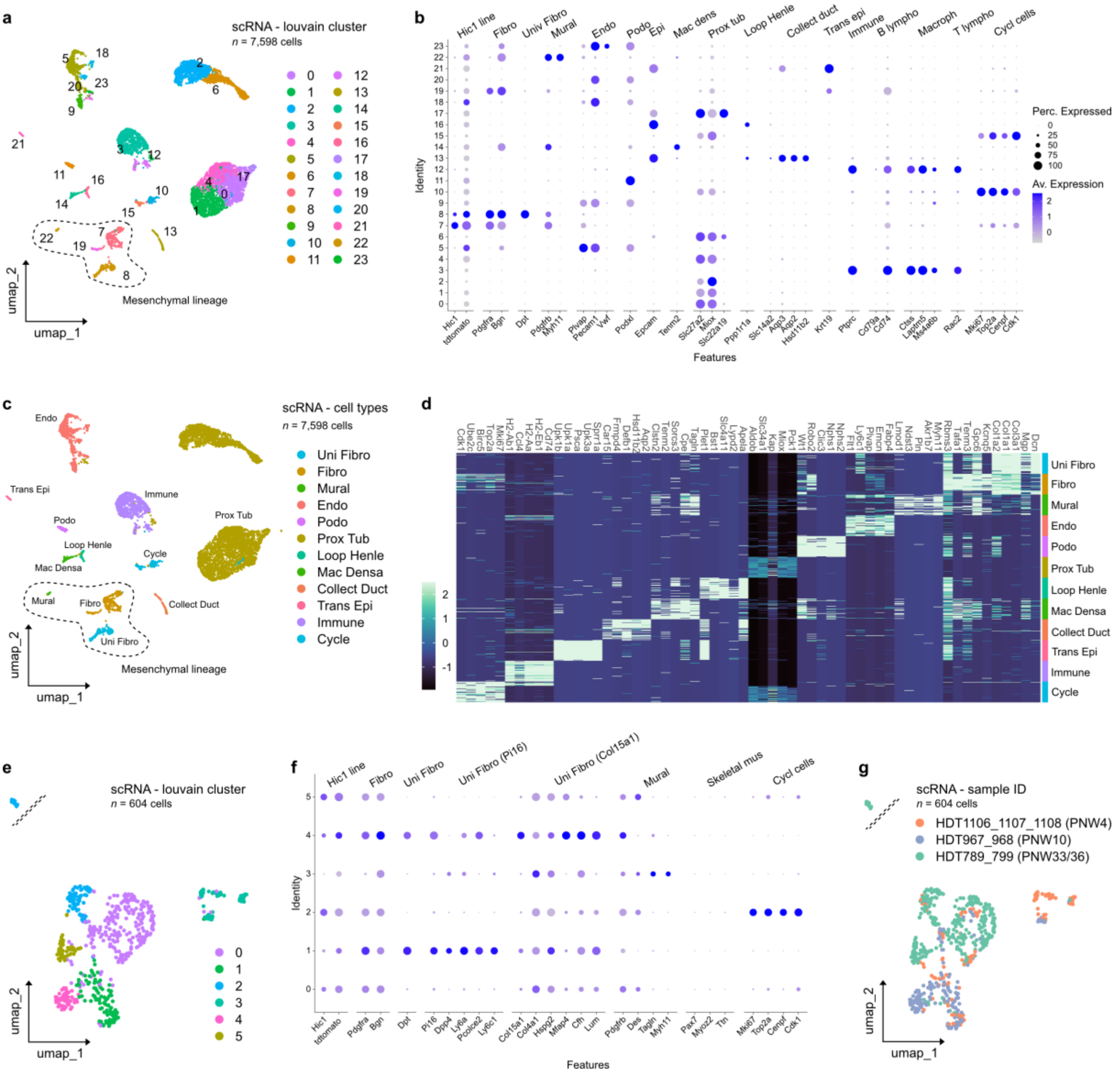


**Extended Data Fig. 3:** **a**, UMAP plot from scRNA-seq data of cells from control mice samples (*n*=3) colored by cluster assignment. Louvain method was used for clustering. **b**, DotPlot showing expression of kidney cell type markers, as well as broad fibroblast and mural cell type markers in identified clusters used for cell type annotation [24-27]. **c**, UMAP plot from scRNA-seq data of cells from control mice samples (*n*=3) colored by assigned cell types, which are annotated based on transcripts representative of different lineages. **d**, Heatmap of top 5 signature defining transcripts, based on cell type annotation. **e**, UMAP plot from scRNA-seq data (*n*=3) of *tdtomato*+ mesenchymal stromal cells colored by cluster assignement. Louvain method was used for clustering (relating to Fig. 2b). **b**, DotPlot showing expression of broad fibroblast and mural cell type markers[25-27] in identified clusters used for cell type annotation. **c**, UMAP plot from scRNA-seq data (*n*=3) of *tdtomato*+ mesenchymal stromal cells colored by sample ID. Louvain method was used for clustering (relating to Fig. 2b).

**Extended Data Fig. 4**

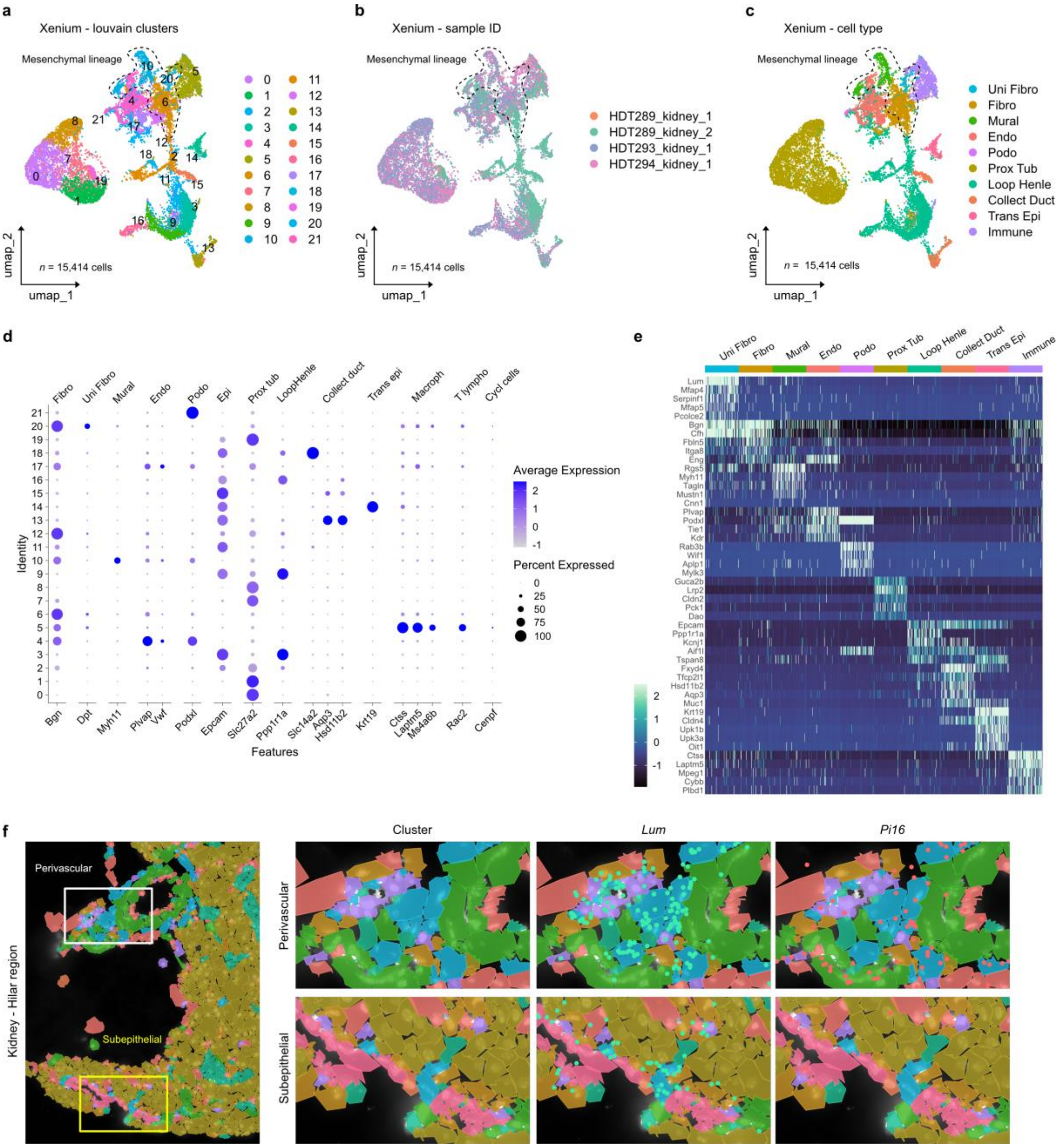


**Extended Data Fig. 4: a-c**, UMAP plot from targeted spatial transcriptomics data (10X Genomics, Xenium, 379 genes) of cells from control mice samples (*n*=4) colored by (**a**) cluster assignment, (**b**) sample ID, and (**c**) assigned cell types, which are annotated based on transcripts representative of different lineages (relating to clusters shown in Fig. 2f-h). Louvain method was used for clustering. **d**, DotPlot showing expression of broad fibroblast and mural cell type markers in identified clusters used for cell type/state annotation [25-27]. **e**, Heatmap of top 5 signature defining transcripts, based on cell type annotation (relating to clusters shown in Fig. 2f-h). **f**, Mapping of clusters identified from targeted spatial transcriptomics to the hilar region of kidney including close-up of the perivascular and subepithelial niche. Coloring of cells corresponds to clusters identified in panel **c**. Expression of universal fibroblast marker *Lum* (identified in panel e) and Uni Fibro *Pi16*$^{high}$ marker (identified in Fig. 2d) *Pi16* is shown.

### Extended Data Fig. 5

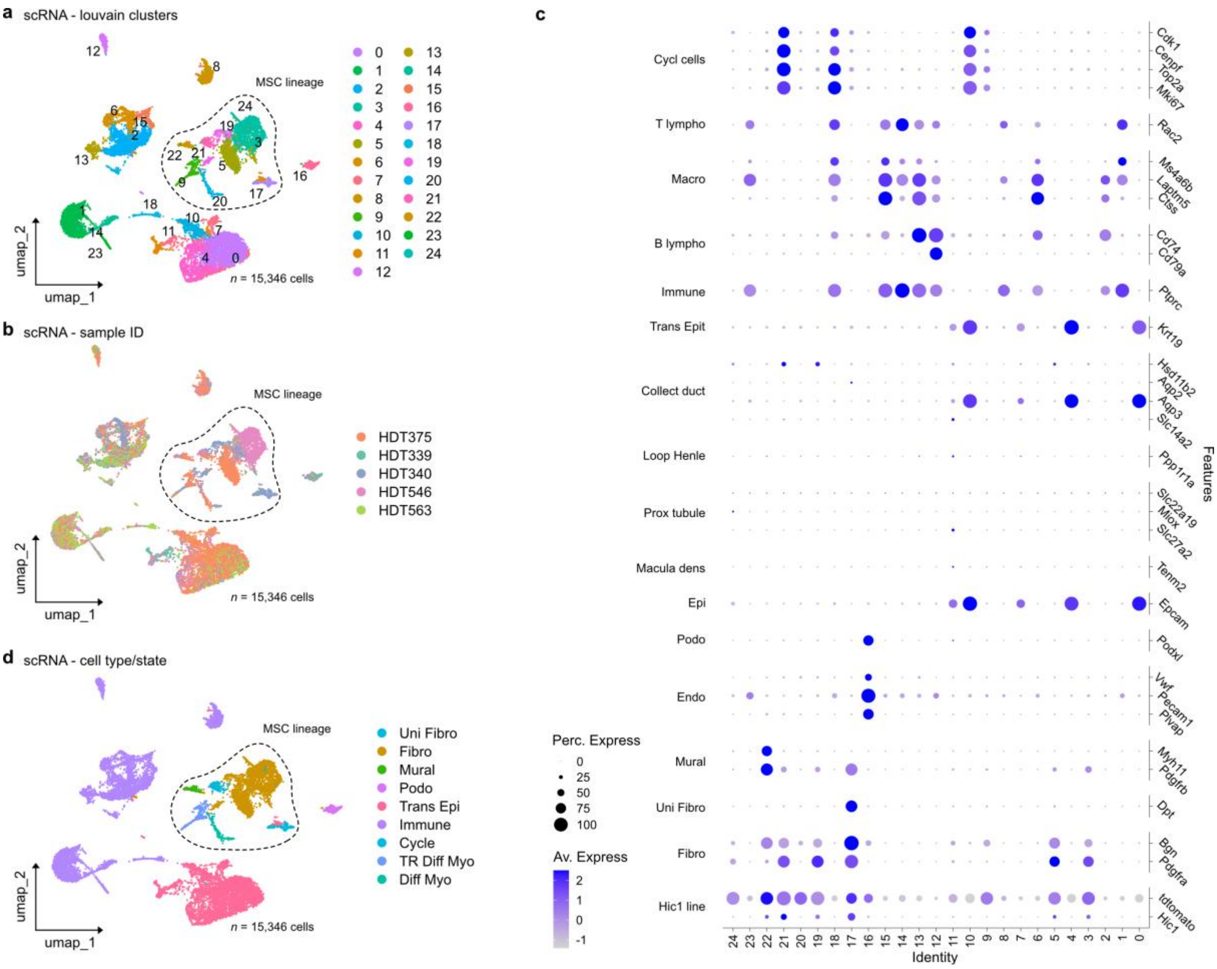


**Extended Data Fig. 5: a**, **b**, and **d**, UMAP plot from scRNA-seq data of cells from HDT tumor samples (*n*=4) colored by (**a**) cluster assignment, (**b**) sample ID, and (**d**) assigned cell type, which are annotated based on transcripts representative of different lineages. Louvain method was used for clustering. **c**, DotPlot showing expression of kidney cell type markers, as well as broad fibroblast and mural cell type markers in identified clusters used for cell type/state annotation[24-27].

**Extended Data Fig. 6**

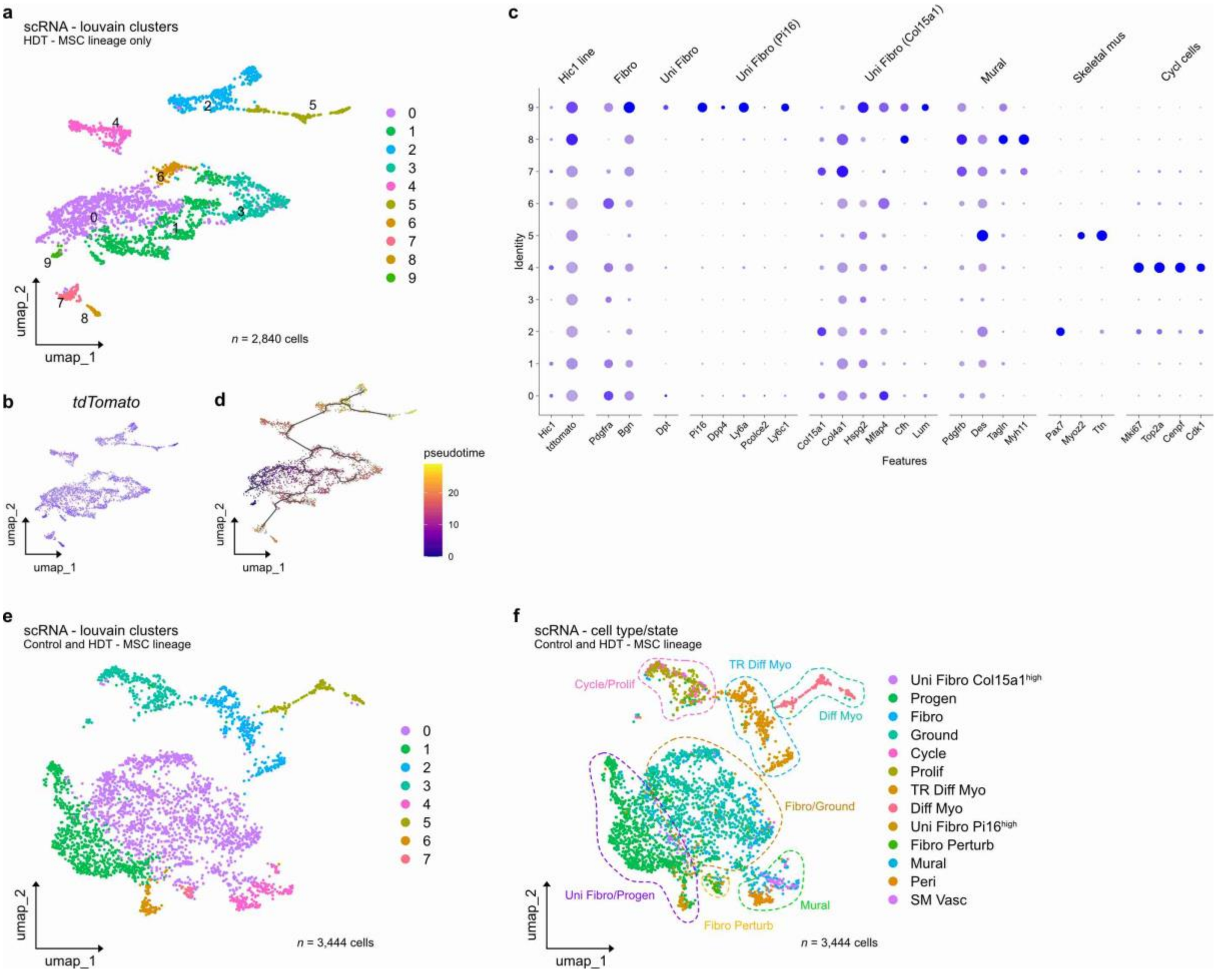


**Extended Data Fig. 6: a**, UMAP plot from scRNA-seq data of cells from HDT tumor samples (*n*=4) colored by cluster assignment. Louvain method was used for clustering. **b**, UMAP plots depicting expression of *tdTomato*. **c**, DotPlot showing expression of broad fibroblast and mural cell type markers in identified clusters used for cell type/state annotation[25-27]. **d**, Trajectory analysis with origin set in Progen near cells with highest *Dpt* expression. Cells are ordered according to pseudotime. **e** and **f**, UMAP plot from combined scRNA-seq data of cells from control mice samples (*n*=3) and cells from HDT tumor samples (*n*=4) colored by (**e**) cluster assignment, and (**f**) cell type assigned during individual work-up of scRNA data (relating to Fig. 2b and Fig. 3b). Louvain method was used for clustering. Broad transcriptional groups, which include cells from either both control and HDT samples or only one of these groups, are indicated by dashed circles.

### Extended Data Fig. 7

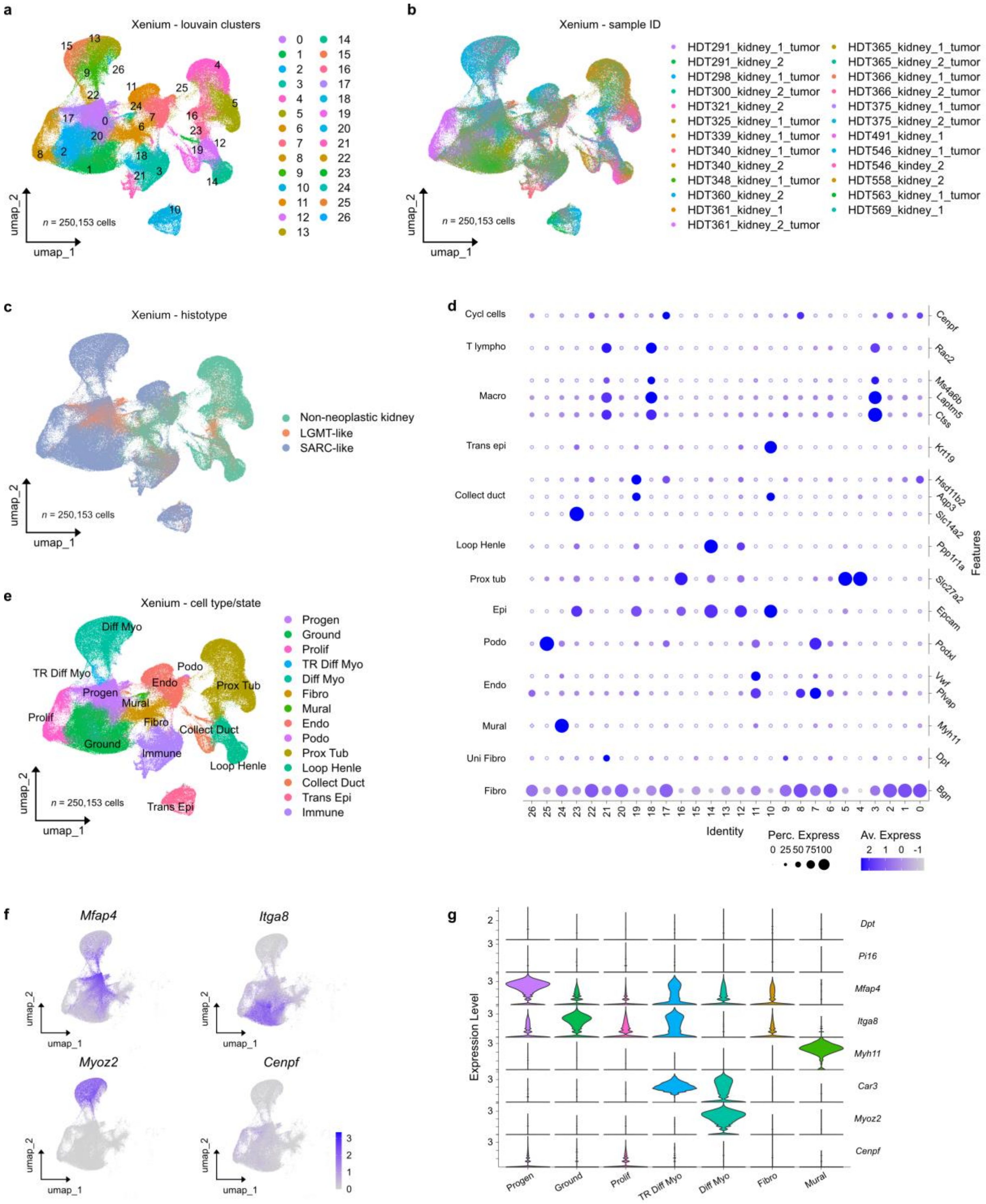


**Extended Data Fig. 7: a**-**c**, and **e**, UMAP plot from targeted spatial transcriptomics data (10X Genomics, Xenium, 379 genes) of cells from HDT mice samples (*n*=25) colored by (**a**) cluster assignment, (**b**) sample ID, and (**c**) histotype, and (**e**) assigned cell types, which are annotated based on transcripts representative of different lineages. **d**, DotPlot showing expression of kidney cell type markers, as well as broad fibroblast and mural cell type markers in identified clusters and used for cell type/state annotation[24-27]. **f** and **g**,

UMAP plots and violin plots of key markers identified in HDT tumor samples (relating to Fig. 3c).

**Extended Data Fig. 8**

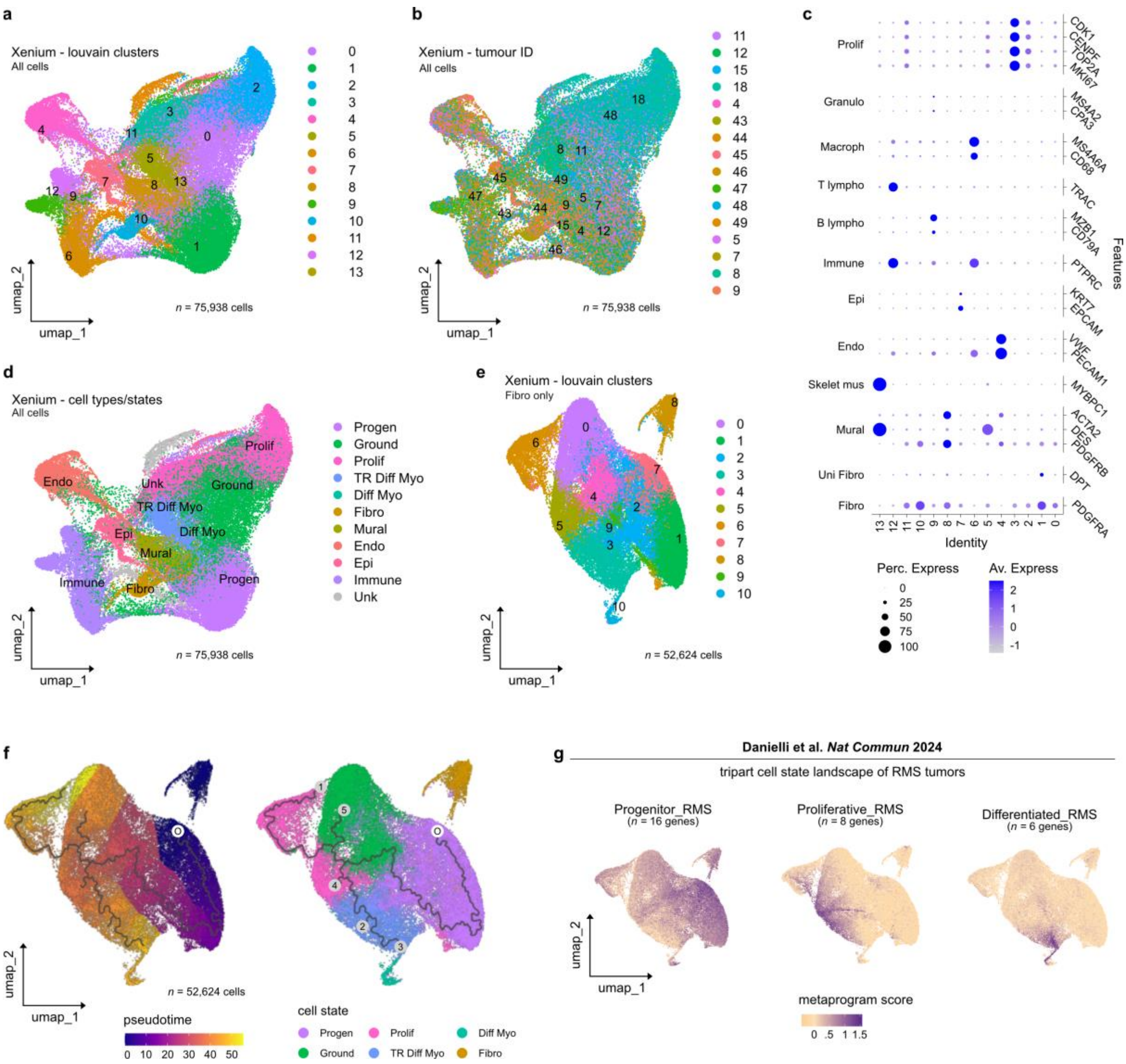

**Extended Data Fig. 8: a**, **b**, and **d,** UMAP plot from targeted spatial transcriptomics data (10X Genomics, Xenium, 377 genes) of cells from patient samples (*n*=16) colored by (**a**) cluster assignment, (**b**) tumor ID, and (**d**) assigned cell types/states, which are annotated based on transcripts representative of different lineages. **c**, DotPlot showing expression of broad cell type and fibroblast markers in identified clusters used for aiding cell type/state annotation[27,32]. **f,** Trajectory analysis of Xenium spatial transcriptomic data with origin set in Progen with high *DPT* expression. UMAP colored by pseudotime (left panel) and cell type/state (right panel). O indicates origin, trajectories are numbered. **g,** Projection of three major RMS tumor metaprograms[33] onto identifies cell states/types of

DICER1-associated mesenchymal tumors using a subset of metaprogram markers present on the targeted panel (see Source Data for Extended Data Fig. 8g).
